# Three-dimensional multi-target super-resolution microscopy of cells using Metal-Induced Energy Transfer and DNA-PAINT

**DOI:** 10.1101/2024.04.02.587536

**Authors:** Nazar Oleksiievets, Nikolaos Mougios, Samrat Basak, Daniel C. Jans, Lara Hauke, Jan Christoph Thiele, Stefan Jakobs, Felipe Opazo, Jörg Enderlein, Roman Tsukanov

## Abstract

Achieving nanometer precision in 3D remains a major challenge in super-resolution microscopy. DNA-PAINT offers excellent lateral resolution and versatile multiplexing, but its axial localization precision is typically 3–5 times poorer, limiting quantitative 3D imaging. Here, we present MIET-PAINT, which combines DNA-PAINT with Metal-Induced Energy Transfer (MIET) to overcome this limitation. We implement MIET-PAINT on both wide-field fluorescence lifetime and confocal TCSPC platforms. Wide-field MIET-PAINT enables robust, multiplexed imaging of focal adhesion proteins and actin in fixed cells. To address the lateral resolution limits of lifetime cameras, we further developed confocal MIET-PAINT, which leverages optical background rejection and high-efficiency SPAD detection. This modality achieves ∼12 nm lateral precision and resolves the 3D architecture of microtubules, vimentin, and actin with high fidelity. MIET-PAINT thus unites nanometer-scale axial accuracy with the multiplexing versatility of DNA-PAINT, establishing a powerful tool for quantitative 3D cell biology.

## INTRODUCTION

Multi-target super-resolution imaging provides powerful insights into the molecular organization and function of complex biological systems, enabling mapping of protein networks, visualization of cytoskeletal architecture, and detailed studies of subcellular structures such as synapses ^1,2^ and focal adhesions^3^. Single-molecule localization microscopy (SMLM)^4,5^ has become a cornerstone in this field due to its ability to break the diffraction limit using relatively simple optical implementations. Among SMLM methods, DNA-PAINT (DNA-based Point Accumulation for Imaging in Nanoscale Topography)^6,7^ is particularly attractive for multiplexed imaging. It achieves nanometer-scale lateral localization precision by using short, fluorescently labeled “imager” strands that transiently bind to complementary “docking” strands on the target. Continuous replenishment of fluorophores circumvents photobleaching, enabling acquisition of densely labeled targets. Orthogonal DNA sequences allow multi-target imaging either sequentially through buffer exchange (Exchange-PAINT)^8,9^ or in parallel using spectral^10,11^, kinetic barcoding^12^, or fluorescence lifetime-based^13,14^ separation. However, cellular Exchange-PAINT imaging can be complicated due to sticky and dense environment inside a cell. Furthermore, the axial resolution of DNA-PAINT typically remains three-fold worse than its lateral precision^5^.

In the realm of SRM, achieving high axial resolution remains a persistent challenge. Several strategies have been developed to address this limitation. Among the most widely used is astigmatic imaging^15^, which is technically simple, broadly compatible with commercial and custom microscopes, and routinely achieves ∼30–60 nm axial precision. Other non-interferometric approaches—such as biplane imaging^16^, variable-angle TIRF^17^, and photometric methods like SIMPLER^18^—extend the axial localization range or improve precision without major hardware modifications, but generally remain in the tens-of-nanometers regime. In contrast, 3D MINFLUX^19^ and interferometric techniques such as iPALM^20^, 4Pi single-molecule detection^21^ achieve true nanometer-scale axial precision by employing dual-objective interferometry or structured excitation schemes. However, these methods require interferometric stability on the nanometer scale and highly specialized optical setups, limiting their widespread adoption. Near-field approaches such as supercritical angle fluorescence (SAF)^22^ and Metal-Induced Energy Transfer (MIET)^23^ provide an attractive alternative, achieving nanometer axial resolution with simpler instrumentation. MIET microscopy offers a compelling alternative, delivering nanometer-scale axial localization over distances up to ∼200 nm from a metal-coated surface using only a fluorescence lifetime information^24^. MIET exploits the steep, distance-dependent quenching of fluorophore emission near a metallic film, which shortens fluorescence lifetime in a predictable manner^25^. This dependency enables precise height determination while maintaining fluorophore photostability. MIET can be implemented on both wide-field^26^ and confocal^27^ fluorescence lifetime microscopes, and has been applied to study a variety of biological systems, such as imaging blood platelet spreading and adhesion^28^, actin cytoskeleton reorganization during epithelial-to-mesenchymal transformation^29^, measuring nuclear envelope inter-bilayer distances^30^, and elucidating the architecture of focal adhesions^31,32^. In addition, single-molecule MIET (smMIET) applications has been demonstrated^26,33,34^. For sub-nanometer axial precision, graphene-induced energy transfer (GIET) offers a steep lifetime–distance dependence, enabling distance measurements up to ∼25 nm above the graphene surface^35–38^. Its precision makes GIET particularly valuable in membrane biophysics and it has been combined with DNA-PAINT for enhanced resolution imaging^39^.

The strengths of MIET and DNA-PAINT are highly complementary: MIET provides unmatched axial resolution, while DNA-PAINT offers virtually unlimited photon budget, flexible multiplexing, and high lateral precision. Here, we introduce MIET-PAINT, which integrates these advantages and demonstrate its implementation on both wide-field fluorescence lifetime imaging and confocal TCSPC-based SMLM platforms. Wide-field MIET-PAINT delivers high-throughput, large-area multiplexed imaging, validated using spacers and applied to mapping the vertical organization of focal adhesion proteins and actin filaments. Confocal MIET-PAINT leverages the high quantum efficiency of SPAD detectors and optical background rejection to achieve improved lateral precision (∼15 nm) in small regions of interest, enabling detailed 3D mapping of cytoskeletal filaments—microtubules, vimentin, and actin. Together, these complementary implementations establish MIET-PAINT as a versatile, accessible platform for multi-target, quantitative 3D super-resolution imaging across diverse biological systems.

The focal adhesion complex (FAC) is a key structure in cell–extracellular matrix (ECM) adhesion, integrating force transduction, cytoskeletal regulation, and tension-dependent signaling^40^. Comprising hundreds of proteins arranged in vertical layers, FACs act as mechano-sensing hubs that extend up to ∼200 nm above the substrate^3,41^. Near-field microscopy methods such as supercritical angle fluorescence (SAF)^22^ and MIET^23^ are ideally suited for resolving their nanoscale architecture, providing insights into adhesion maturation, turnover, and responses to mechanical or biochemical perturbations^42^. We have previously used MIET to map the vertical arrangement of actin, vinculin, and stress fibers in fixed cells and to demonstrate its feasibility for live-cell imaging^31,43^.

Here, we apply multi-target MIET-PAINT to image three representative FAC components spanning distinct layers: paxillin (integrin-signaling layer), zyxin (actin-regulatory layer), and actin filaments (force-transmission network)^3,44^. Paxillin, particularly its phosphorylated form pPax-Y118, is an early adhesion marker enriched in mature FACs, where it modulates integrin signaling and adhesion stability^42,44–46^. Zyxin localizes to both the actin-regulatory layer and stress fibers, contributing to actin repair and reinforcement in a tension-dependent manner^42,45,47–50^. Actin stress fibers form the principal tracks for transmitting forces between the ECM and cytoskeleton, making them essential for mechanotransduction studies^40,44,48^. Beyond FACs, the cytoskeleton—comprising actin filaments, intermediate filaments such as vimentin, and microtubules—maintains cell shape, organizes organelle positioning, supports intracellular transport, and coordinates mechanical responses^51^. Actin provides structural support and contractile force, vimentin imparts mechanical resilience, and microtubules enable long-range transport and establish cell polarity^52^. Mapping their nanoscale 3D organization is critical for understanding how cells integrate structure and mechanics^53^.

## RESULTS

### Principle and validation of MIET-PAINT

To verify the efficacy of the MIET-PAINT technique, we designed a validation sample for imaging on top of SiO2 layers of varying known thicknesses. This validation method, previously successful for calibrating MIET with stationary Atto 655 fluorophores^26^, has now been adapted for MIET-PAINT imaging by passivating the surface with docking strands, as illustrated in Figure 1a. We prepared gold slides with SiO_2_ spacers of different thicknesses, achieving sub-nanometer precision in their preparation. Biotin-avidin chemistry was employed to specifically bind biotinylated docking strands to the SiO_2_ surface, adding an additional ∼12 nm height from the biotin-avidin complex (“immobilization sandwich”) to the fluorophores’ absolute height above the gold layer^34^.

**Figure 1.**
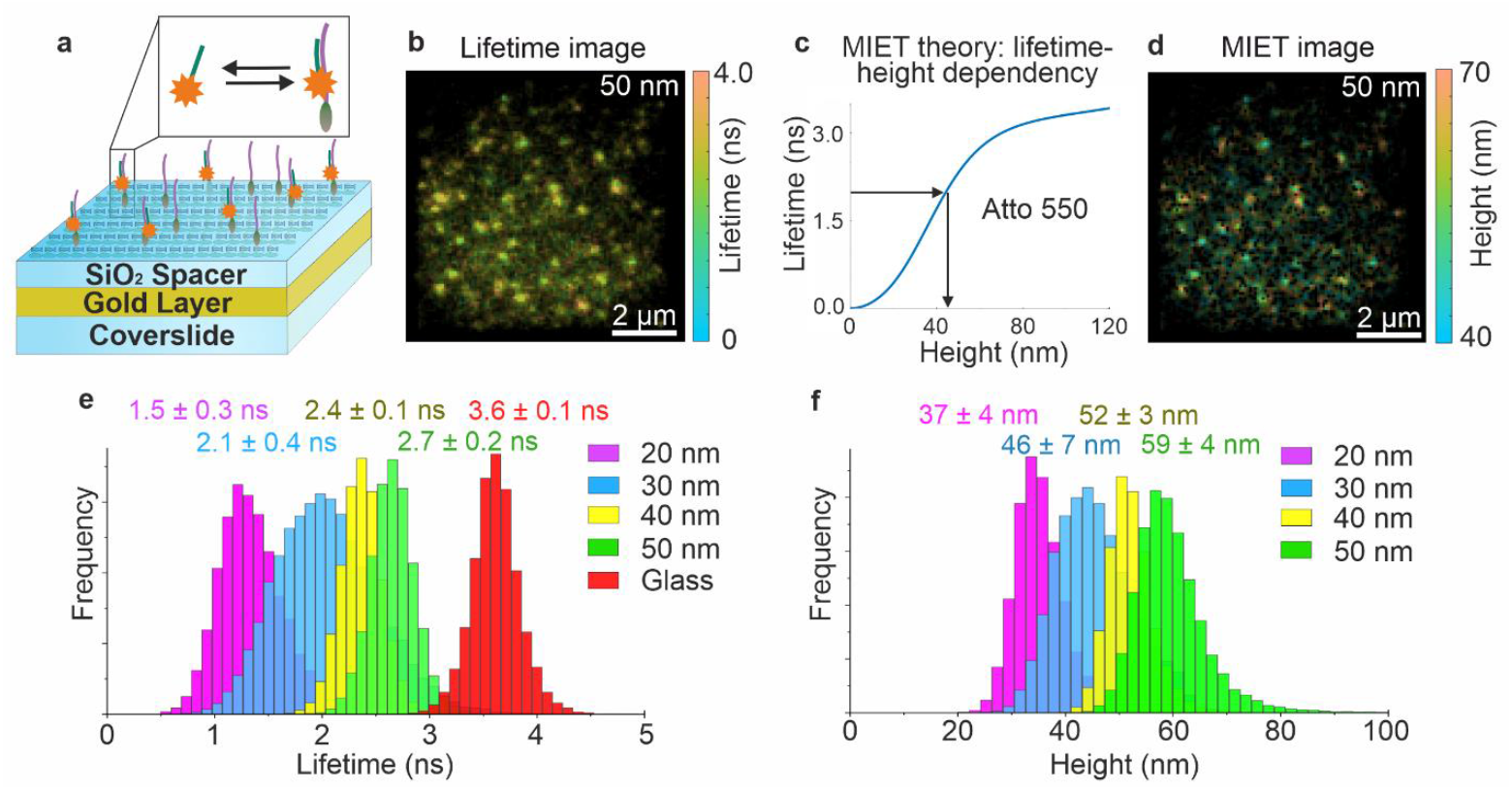
The principle and validation of MIET-PAINT technique. (a) MIET-PAINT validation experiment design: gold glasses covered with SiO2 spacers with different thicknesses are functionalized with docking strands via biotin-avidin chemistry; imager strand present in solution transiently binds to the complementary docking strand resulting in binding events. (b) Exemplary single-molecule MIET-PAINT lifetime image for 50 nm SiO2 spacer (c) MIET theory: calculated lifetime-distance dependency for Atto 550 fluorophore attached to DNA. (d) Exemplary single-molecule MIET-PAINT height image obtained from (c). (e) Lifetime and (f) height histograms for different SiO2 spacers thicknesses. The extra height of 12 nm is present in the experimental value due to the additional thickness of the immobilization layer. The average values and errors are displayed in colors that match those of the histograms.

During the experiments, imager strands were introduced into the chamber, and MIET-PAINT imaging was conducted using our wide-field fluorescence lifetime imaging microscopy (FLIM) optical setup, detailed in Figure S1 of the Supplementary Information (SI).

To expedite DNA-PAINT acquisition, we utilized an optimized imager strand (R4-Atto 550) that avoids the formation of secondary structures, thereby accelerating the binding rate. This development was introduced by the Jungmann lab^54^. MIET-PAINT imaging was performed on SiO_2_ spacers with thicknesses of 20, 30, 40, and 50 nm. Exemplary lifetime images are shown in Figure 1b, with the theoretical lifetime-distance dependency (MIET curve) used to convert lifetime values into heights depicted in Figure 1c, and an exemplary 3D MIET-PAINT image presented in Figure 1d. For each set of measurements, we generated a lifetime histogram and a corresponding height histogram, as displayed in Figures 1e-f and S4. Upon comparing the experimentally measured heights with the designed height values, we observed complete agreement, as documented in Table S1 of the SI. It’s important to note that, for accurate comparison between measured and designed heights, an additional ∼12 nm must be considered to account for the thickness of the biotin-avidin immobilization layer. This comprehensive validation approach effectively demonstrates MIET-PAINT’s precision and reliability for measuring fluorophore heights with nanometer accuracy within the sensitive MIET range of 100 nm above a surface.

### Multi-target MIET-PAINT imaging

#### Wide-field MIET-PAINT imaging of Focal Adhesion Complex

We applied wide-field MIET-PAINT to image paxillin (pPax-Y118), zyxin, and actin stress fibers in genome-edited Zyxin-rsEGFP2 U2OS cells. To decrease target-fluorophore distance (linkage error), targets were labeled with nanobodies^9^ coupled to DNA docking strands or LifeAct^55^, and imaged sequentially using Exchange-PAINT with optimized imagers (R1, R4)^54^ and LifeAct (Figure 2a–c). The sequential workflow enabled chromatic-aberration-free 3D imaging with a single spectral channel. For each target, lateral coordinates of emitters were determined from single-molecule localizations, and axial positions were extracted from fluorescence lifetimes via the MIET calibration curve (Figures 2d–f), as described in Methods section. From these data, height profiles were generated and directly compared across targets, allowing us to assess the vertical positioning of zyxin, paxillin, and actin within the same focal adhesion (Figures 2g–h, S6, S7). This analysis uncovered a clear vertical stratification of focal adhesion proteins. Phosphorylated paxillin localized at 53 ± 12 nm above the substrate, zyxin at 74 ± 18 nm, and actin filaments at > 100 nm, as visualized in Figure S8 and Supplementary Videos. Zyxin showed partial colocalization with actin stress fibers, consistent with its recruitment to tension-bearing structures, and cross-correlation analysis (Pearson coefficient) demonstrated strong spatial correlation of zyxin with both paxillin and actin (Figure S12). Actin fibers inclined at an average of 0.7°, in agreement with previous MIET measurements. The average localization precisions were 22 nm lateral and 5 nm axial (Table S2). These results confirm MIET-PAINT’s capacity for quantitative, multi-target 3D mapping of nanoscale protein organization within focal adhesions.

**Figure 2.**
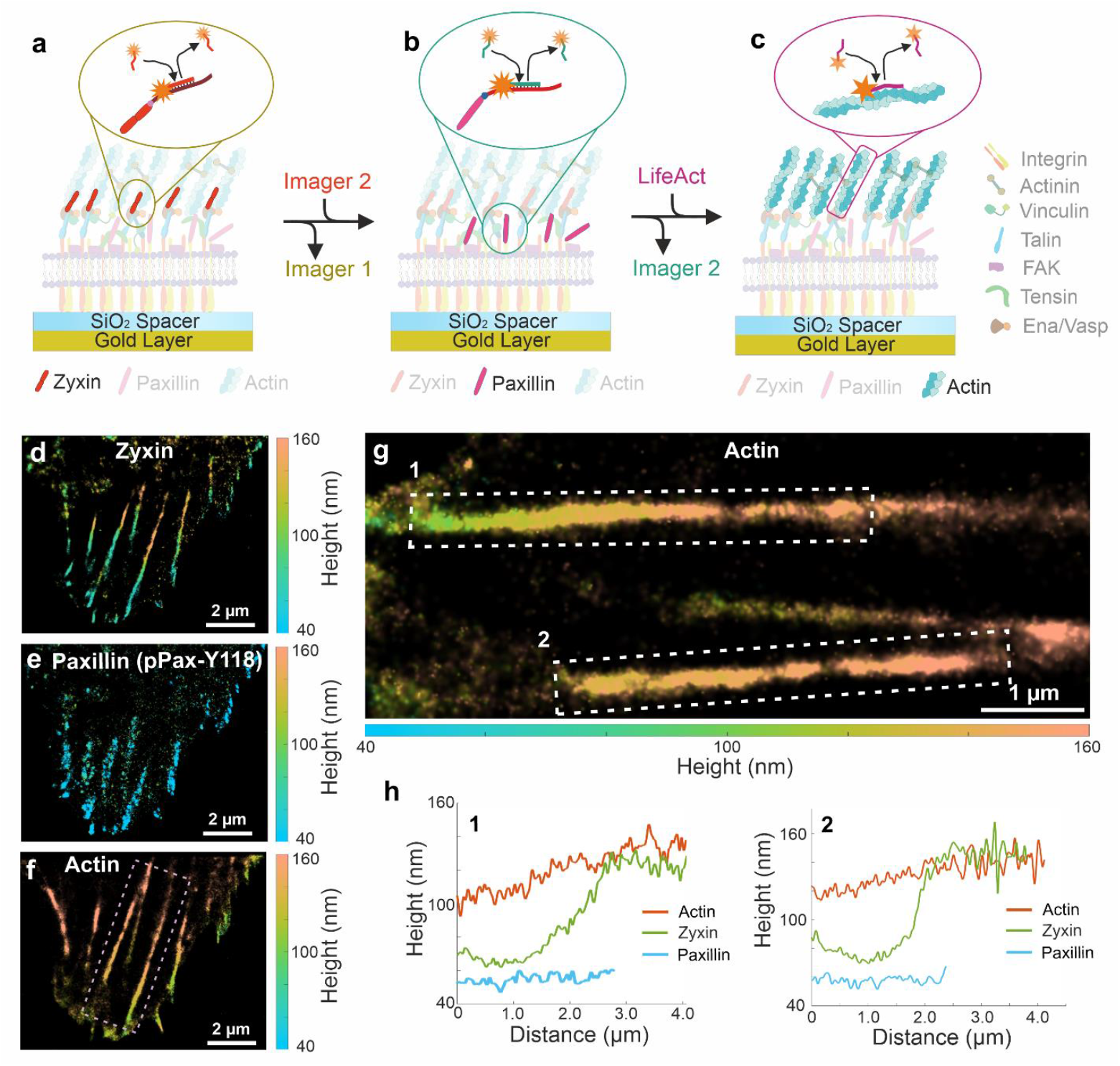
Wide-field MIET-PAINT imaging of FAC and actin stress fibers. (a-c) Schematic of multi-target MIET-PAINT experiment demonstrated for the imaging of Focal Adhesion Complex proteins (zyxin and paxillin) and actin stress fibers. The targets are imaged sequentially, while each imaging cycle is performed with different imagers (or LifeAct in case of actin imaging). In between the imaging cycle, the previous imager is washed out and the next imager is introduced. MIET-PAINT images of zyxin (d), paxillin (pPax-Y118) (e), and actin (f) of the same cell. All scale bars are 2 µm. (g) Zoom-in region of actin image shown in (f) in pink dashed rectangle. (h) Line height profiles of zyxin, paxillin, and actin along the stress fibers. The numbers depict the selection of two stress fibers as shown in (g) in white dashed rectangles.

#### Confocal MIET-PAINT imaging of cytoskeleton

To extend MIET-PAINT’s versatility, we developed a confocal variant using laser-scanning fluorescence lifetime detection compatible with SMLM^27^. This approach achieved improved localization precisions (12.6 nm lateral, 5.7 nm axial, Table S3.) owing to high detector quantum efficiency and out-of-focus background rejection, making it well suited for MIET-PAINT imaging of small regions of interest (≤ 20 × 20 µm^2^). Sequential imaging of microtubules, vimentin, and actin in fixed mammalian cells revealed their 3D organization and spatial relationships (Figure 3a-c, S9, S10, S11). Quantitative co-localization analysis (Pearson correlation) showed substantial overlap between vimentin–microtubules and actin– microtubules, but low colocalization between actin and vimentin (Figures 3d, S12), consistent with their distinct but interconnected roles in cytoskeletal architecture. Representative actin height profiles are shown in Figure 3e–f. These results demonstrate that confocal MIET-PAINT complements wide-field implementation by providing high-precision 3D imaging in targeted cellular regions.

**Figure 3.**
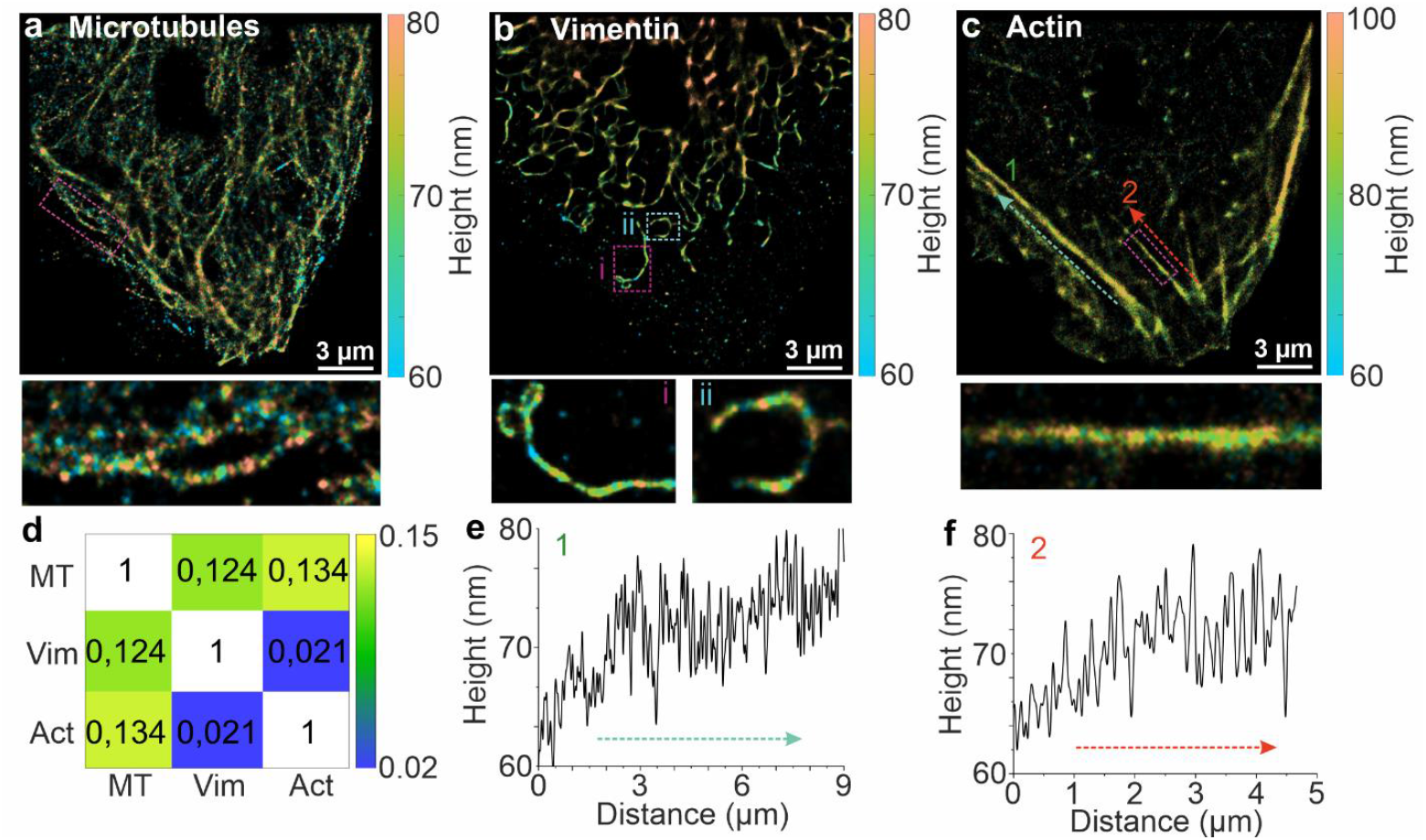
Confocal MIET-PAINT imaging of cytoskeleton. Sequential MIET-PAINT imaging of microtubules (a), vimentin (b) and actin (c). Between the imaging cycles, through washing was employed to avoid the crosstalk between the targets. All scale bars are 3 µm. Below are the Zoom-in regions of images shown in (a-c) in pink dashed rectangles. (d) A heat map of the Pearson’s r-value. (e,f) Line height profiles of actin along the stress fibers. The numbers depict the selection of two stress fibers as shown in (c) marked by cyan and red dashed lines.

## DISCUSSION

MIET-PAINT combines the high axial precision of Metal-Induced Energy Transfer (MIET) with the multiplexing capacity and versatility of DNA-PAINT, enabling isotropic nanometer-scale 3D localization (∼25 nm lateral, ∼5 nm axial). This integration overcomes the limitations of previously introduced MIET-dSTORM, which—while capable of precise 3D localization—are constrained by limited multiplexing, and reliance on a narrow palette of well-blinking dyes. In MIET-dSTORM, photon budget is finite, acquisition time is limited, and dense molecular sampling is difficult to sustain, particularly in multi-target experiments. DNA-PAINT, by contrast, provides an essentially unlimited photon budget, dense sampling, and multiplexing via orthogonal DNA sequences. MIET-PAINT leverages these complementary strengths, enabling long, high-density, and multi-target 3D acquisitions without compromise.

Our wide-field MIET-PAINT implementation proved robust and high-throughput, enabling detailed 3D mapping of focal adhesion complexes (FACs), including the axial co-localization of paxillin, zyxin, and actin stress fibers on a single adhesion level. High axial localization precision was achieved using a LINCam TCSPC camera, while HILO and TIRF illumination improved contrast. Lateral localization was limited to ∼25 nm due to the modest quantum efficiency of the LINCam S20 detector. Ongoing developments in wide-field FLIM—such as next-generation lifetime cameras with improved quantum efficiency—are expected to further enhance lateral and axial resolution^56^.

To further enhance lateral resolution, we developed a confocal MIET-PAINT variant on a laser-scanning SMLM platform with a SPAD detector, achieving ∼12 nm lateral precision in ROIs of 20 × 20 µm^2^ while maintaining comparable axial resolution. The confocal pinhole provided strong background rejection, making the approach ideal for small-area, high-precision imaging, albeit at the cost of longer acquisition times and increased setup complexity.

Both MIET-PAINT modalities are compatible with Exchange-PAINT workflows, enabling theoretically unlimited multiplexing with a single fluorophore, eliminating chromatic aberration. Together, wide-field MIET-PAINT offers large-area, high-throughput imaging, while confocal MIET-PAINT excels in small-volume, ultra-precise measurements—forming a versatile toolkit for nanoscale exploration of cellular architecture. Importantly, by enabling multi-target 3D mapping of FAC proteins and cytoskeletal networks with nanometer resolution in the same cell, MIET-PAINT opens new opportunities to dissect how nanoscale spatial relationships reveal cellular adhesion, force transmission, and structural integrity.

## METHODS

### Validation of MIET-PAINT

We designed an experiment for MIET-PAINT validation as follows: the imager binding events were localized on top of SiO_2_ spacers with different precisely determined thicknesses. Previously, such MIET validation strategy has proved to be successful for stationary Atto 655 fluorophores validation^26^. For MIET-PAINT validation, we employed docking strand R4*^54^, sequence (5’→3’): Biotin-ACACACACACACACACACA, for details see Table S2 in the SI. We used the docking strand R4* modified with biotin on its 5’ end and immobilized it onto coverslip surface functionalized with BSA-biotin (A8549, Sigma-Aldrich) and then Neutravidin (31000, Thermo Fisher Scientific). We selected this docking strand as it has been used in the MIET-PAINT imaging of cells. The thickness of the biotin-avidin immobilization layer was reported to be ∼12 nm^34^, and was added to the spacer thickness to get total height value of an emitter above gold surface. The imager R4 had the following sequence (5’→3’): TGTGTGTTT and carried Atto 550 fluorophore on its 3’ end. The imager was diluted to a concentration of ∼0.5 nM in the imaging buffer PBS pH 7.4 with NaCl 500 mM and was injected into the Petri dish (Ibidi 81158, Germany). The imager-docking bindings events were detected and localized, while fluorescence lifetime for each event was extracted from TCSPC curves, see Figure S3 in the SI. Then, the lifetime values were converted into axial positions, resulting in an emitter localization in 3D, see Table S1 in the SI. Series of measurements with different spacer thicknesses above gold layer has been performed using a wide-field FLIM setup, as shown in Figure S1 in the SI.

### Cell culture

We used genome-edited U2OS cell line Zyxin-rsEGFP2 (homozygous), introduced previously in Ref^57^. The cells were cultivated in McCoys 5a medium (16600082, Thermo Fisher Scientific), supplemented with 100 U ml^−1^ penicillin, 100 ug ml^−1^ streptomycin (P0781, Sigma-Aldrich), 1 mM Na-pyruvate (S8636, Sigma-Aldrich) and 10% (v/v) FBS (FBS.S 0615, Bio and SELL) at 37°C, 5% CO_2_.

### Cell sample preparation

The cells were cultured for 1 day on 35 mm glass bottom (No. 1.5) dishes (81158, Ibidi), or, alternatively, in the same dishes but with gold coating. For the latter, following layers were used: 10 nm gold and 5 nm SiO_2_. The cells were fixed in pre-warmed 8% formaldehyde in PBS for 10 min. After washing with PBS pH 7.4 the dish was stored at 4°C before imaging.

### Immunostaining of cells

Cells were permeabilized and blocked using 2% bovine serum albumin (BSA) and 0.2% Triton X-100 in PBS for 30 min at room temperature. PBS buffer containing nanobodies coupled to a docking strand (nanobodies concentration in range 20-50 nM) was used to stain the cells. The incubation time varied between 1-2h at room temperature. Finally, cells were washed three times for 15 minutes in total, rinsed with PBS buffer and then post-fixed with 4% PFA for 15 min at room temperature. The remaining aldehydes were quenched by 0.1 M glycine in PBS. Cells were stored in PBS at 4°C. The unconjugated nanobodies FluoTag-Q anti-GFP (NanoTag Biotechnologies GmbH, Germany, Cat. No: N0301) carrying one ectopic cysteine at the C-terminus allow for chemical couplings via a thiol-reactive compound. DNA docking strands (Biomers GmbH, Germany) were functionalized with an azide group at 5’ end. The docking strands were coupled to the nanobodies using a dibenzocyclooctyne (DBCO) crosslinker^9^. FluoTag-Q anti-GFP was coupled to R4* concatenated sequence 5’-TACACACACACACACACACA-3’^54^. To label paxillin we employed the pre-mixing protocol^58^. Recombinant Anti-Paxillin (pPax-Y118) antibody (Abcam, ab32084, concentration 15 nM) and FluoTag-X2 anti-Rabbit IgG (NanoTag Biotechnologies GmbH, N2402, Germany, the concentration of 45 nM) were pre-mixed in the tube and incubated for 30 minutes. FluoTag-X2 anti-Rabbit IgG was coupled to R1* docking sequence 5’-TCCTCCTCCTCCTCCTCCT-3’^54^. The fluorophore-labeled imager strands were purchased from Eurofins Genomics, Germany. The imagers were modified with Atto 550 fluorophore at its 3’ end, see Table S4 for all DNA sequences and its modifications. The imagers were aliquoted in TE buffer (Tris 10 mM, EDTA 1 mM, pH 8.0) at the concentration of 1 µM and stored at -20°C. Prior to the experiment, the strands were thawed and diluted to the final concentrations of 0.5 nM in PBS buffer pH 7.4 with 500 mM NaCl. Imager solution (∼1 mL) was injected into a Petri dish with cells and MIET-PAINT imaging was performed. For actin imaging, we used commercially available LifeAct functionalized with Cy3B (Cambridge Research Biochemicals, crb1130929h, Billingham, UK). The original solution was diluted to a concentration of 1 µM, aliquoted and stored at -20°C. For imaging, we used LifeAct-Cy3B in a concentration of ∼0.5 nM in PBS buffer pH 7.4 with 500 mM NaCl.

Cytoskeleton labeling for confocal MIET-PAINT was performed as follows: in U2OS cell line microtubules were labeled using anti-α-Tubulin mouse IgG1 (Synaptic Systems, Cat. No: 302211, Germany), pre-mixed with FluoTag-X2 anti-Mouse IgG1 nanobody (NanoTag Biotechnologies GmbH, Cat. No: N2005, Germany) coupled to the R3* docking strand with concatenated sequence 5′-CTCTCTCTCTCTCTCTCTC-3′^54^. Vimentin was labeled using anti-Vimentin rabbit antibody (Abcam, ab92547, Germany), pre-mixed with FluoTag-X2 anti-Rabbit IgG nanobody (NanoTag Biotechnologies GmbH, Cat. No: N2402, Germany) coupled to the R6* docking strand with concatenated sequence 5′-AACAACAACAACAACAACAA-3′^54^.

## Data acquisition

### Wide-field MIET-PAINT imaging of Zyxin cells

Wide-field MIET-PAINT was performed on a custom-built setup (Figure S1, SI) equipped with a pulsed super-continuum white light laser (Fianium WhiteLase SC450, NKT Photonics; 20 MHz repetition rate) and a lifetime camera LINCam25 S20 (Photonscore GmbH)^59^. A custom photodiode was used to trigger the lifetime camera. TIRF illumination was employed for optimal image contrast, with ∼10 mW laser power after the fiber (∼0.5 kW/cm^2^ at the sample). Cells were first screened using an emCCD camera to identify those with mature focal adhesions based on Zyxin–rsGFP fluorescence signal (Figure S5), using simultaneous 405 nm (activation) and 488 nm (excitation) illumination. Once selected, imaging switched to the lifetime camera for MIET-PAINT acquisition. The MIET-PAINT data were acquired using Photonscore Capture software, with typical acquisition times of ∼40 min per target (corresponds to 8k frames with integration time of 0.3 s), typically registering on average 100-150k localizations for zyxin and paxillin, and ∼200k localizations for actin. The targets were imaged sequentially using Exchange-PAINT: Zyxin (R4–Atto 550), paxillin pPax–Y118 (R1–Atto 550), and actin (LifeAct–Cy3B). Imager strands were diluted to ∼0.5 nM in imaging buffer (PBS, 500 mM NaCl) and sequentially introduced into the experimental chamber. Between imaging rounds, chamber washing with PBS buffer was performed using at least three times the volume of Petri dish (∼1 mL), until no residual localizations from the previous imaging round were detected.

### Confocal MIET-PAINT imaging of Cytoskeleton

Confocal MIET-PAINT was implemented on a custom-built laser-scanning confocal microscope with time-correlated single-photon counting (TCSPC) detection (Figure S2, SI), compatible with confocal single-molecule localization microscopy^27^. A pulsed white light laser (Koheras SuperK Power, NKT Photonics) provided flexible excitation wavelength selection. Cells were first scanned over an 80 × 80 µm^2^ ROI to locate suitable candidates for imaging. For MIET-PAINT acquisition, the excitation wavelength was set to 560 nm (560/9 nm clean-up filter), with ∼50 µW power at the objective. Emission was filtered using a 560 nm long-pass and 609/62 nm band-pass filter, optimized for Atto 550 and Cy3B detection.

Sequential Exchange-PAINT imaging targeted microtubules (R3–Atto 550), vimentin (R6–Atto 550), and actin (LifeAct–Cy3B). For each target, a higher imager concentration (∼1 nM) was used to locate and focus on the region of interest, followed by dilution to ∼0.2 nM for single-molecule detection. Data were acquired in bi-directional scan mode (pixel size 100 nm, dwell time 2.5 µs) over a 20 × 20 µm^2^ region, recording 50,000 frames per target, imaging time ∼2h/cycle. On average, 700k localizations were detected per imaging cycle for microtubules and vimentin, and 940k localizations for actin. Between imaging rounds, thorough PBS washing was performed using washing volumes at least three times the volume of Petri dish (∼1 mL), until no residual localizations remained. The focal plane was manually kept constant across rounds to maintain axial registration.

### Data analysis of MIET-PAINT experiment

For MIET-PAINT data analysis, we used previously published Matlab-based TrackNTrace analyse package^60^ available online via link: https://github.com/scstein/TrackNTrace. In the current work, Matlab version R2022a was used for TrackNTrace data analysis. To obtain an emitter axial position, experimentally measured lifetime value of an emitter was converted into height using the theoretical MIET model^25^. Using literature values for refractive indices of gold and titanium literature^61^ and given the sample geometry, lifetime-distance dependency (MIET curve) was calculated for the DNA-Atto 550 conjugate, as shown in Figure S2 in the SI. The following parameters were used for MIET curve calculation: emission wavelength center 576 nm, quantum yield of 0.9, and free space lifetime of 3.74 ns. MIET curve for LifeAct-Cy3B was calculated using the following parameters: emission wavelength center 571 nm, quantum yield of 0.9, and free space lifetime of 2.7 ns. We assumed random orientation of emission dipole, as the C6 linker was used between a fluorophore and DNA. As the output format of the LINCam25 lifetime camera does not include division into pixels and time bin, therefore we were free to choose a spatial binning of 8 pixels, which corresponds to a virtual pixel size of 192 nm, and a time bin of 300 ms. The detection and precise sub-pixel localization of emitters was performed using a cross-correlation algorithm and pixel-integrated Gaussian MLE fitting. Non-repeating localizations detected in a single frame were discarded. For each localization, lifetime information has been extracted by tail-fitting of a TCSPC curve with single-exponential function. For the purpose, TCSPC curve was cut 0.1 ns after the maximum, and the remaining decay was fitted with a mono-exponential function using a maximum likelihood estimator (MLE). A dataset was corrected for a lateral mechanical drift using redundant cross-correlation algorithm^62^ and, finally, a MIET-PAINT image was reconstructed. Following filters were applied to improve image quality: (1) localizations with more than 50 photons, (2) PSF sigma less than 1.8 px and (3) only lifetime values in the range from 0.5 ns to 5.0 ns were taken into account. Height profiles were generated by integrating axial localization data along the axis of interest within a typically 300 nm-wide band. For each position along the axis, heights of localizations in perpendicular cross-sections were averaged, and the resulting values were plotted as a cumulative height profile.

### Corresponding authors

E-mail:(R.T.). Phone: +49 551 39 26911. E-mail:(J.E.).

Phone: +49 551 39 26908.

## Notes

FO is a shareholder of NanoTag Biotechnologies GmbH. All other authors declare no competing interests.

## Supplementary information

Wide-field FLIM optical setup for MIET-PAINT, Theoretical lifetime-height dependency - MIET curves, Buffer solutions, Validation of MIET-PAINT, Fluorescence lifetime decay curves, DNA sequences of imager and docking strands, Additional MIET-PAINT images, Quantitative characteristics of MIET-PAINT images.

## Acknowledgment

The authors are grateful to Photonscore GmbH and personally to Yury Prokazov and André Weber for their support, valuable advice and contribution to the data analysis package. We thank Ingo Gregor for help with 3D data visualization. We are grateful to Anna Chizhik for the preparation of gold-coated coverslips and Petri dishes. The authors thank Sebastian Isbaner for help during the initial stage of the project.

