## Supplementary Information for "Three-dimensional multi-target super-resolution microscopy of cells using Metal-Induced Energy Transfer and DNA-PAINT"

### Wide-field MIET-PAINT

**Optical setup:** wide-field measurements were performed using a custom-built optical setup equipped with lifetime camera, see Figure S1. Pulsed super-continuum white light laser (Fianium WhiteLase SC450, NKT Photonics) with a fixed pulse repetition rate of 20 MHz was employed for excitation. The lifetime-based camera (LINCam25, Photonscore) was optically triggered using a custom-built photodiode (PD)[1]. A clean-up filters (CUF) (BrightLine HC 563/9, Semrock; ZET 488/10, Chroma) were positioned in front of the white light laser output to excite in different wavelengths. Neutral density filters (NE10A-A and NE20A-A, Thorlabs), as well as the variable neutral density filter (ND) (NDC-50C-4-A, Thorlabs) were used to adjust the laser excitation power. The laser beam was coupled into a single-mode optical fiber (SMF) (P1-460B-FC-2, Thorlabs) with a typical coupling efficiency of 30%. After exiting the optical fiber, the collimated laser beam was expanded by a factor of 3.6 ×. The typical excitation intensity at the sample was 10 − 20 W/cm2. The laser beam was focused onto the back focal plane of the TIRF objective (UAPON 100× oil, 1.49 NA, Olympus) using achromatic lens (L1) (AC508-180-AB, Thorlabs). Mechanical shifting of the beam with respect to the optical axis was done through a translation stage (TS) (LNR25/M, Thorlabs) for switching between EPI, HILO, and TIR illumination schemes. The smooth lateral sample positioning was achieved by using a high-performance two-axis linear stage (M-406, Newport). In addition, an independent one-dimensional translation stage (LNR25/M, Thorlabs) together with a differential micrometer screw (DRV3, Thorlabs) was used to move the objective along the optical axis for focusing. The spectral separation of the collected fluorescence light from the excitation pathway was achieved using a multi-band dichroic mirror DM (Di03-R405/488/532/635, Semrock), which direct the fluorescence light towards the tube lens L2 (AC254-200-A-ML, Thorlabs). The field of view was physically limited in the emission path by an adjustable slit aperture (SP60, OWIS) positioned in the image plane in order to image a region with a uniform excitation intensity distribution and also to limit the photon flux reaching the lifetime camera. Lenses L3 (AC254-100-A, Thorlabs) and L4 (AC508-150-A-ML, Thorlabs) re-imaged the emitted fluorescence light form the slit onto an emCCD camera (iXon Ultra 897, Andor). Alternatively, lens L5 (AC508-250-A-MC, Thorlabs) re-imaged the light onto the lifetime camera (LINCam25, Photonscore). The switching between the two cameras was achieved by using a dielectric mirror (BB1-E02, Thorlabs) positioned on a magnetic base plate MB (KB50/M, Thorlabs) with removable top. A band-pass filters BP (BrightLine HC 525/45 Semrock; BrightLine HC 609/54, Semrock; BrightLine FF 962/40, Semrock) were used to block the scattered excitation light. The total magnification of the system on emCCD camera was 166.6×, resulting in an effective pixel size in the sample space of 103.5 nm. The total magnification for the lifetime camera was 222×, and divided into 512 × 512 pixels with the effective pixel size in sample space of 191.6 nm. All experiments were done at 23 ± 1◦C. This was crucial for the mechanical stability of the optical setup.

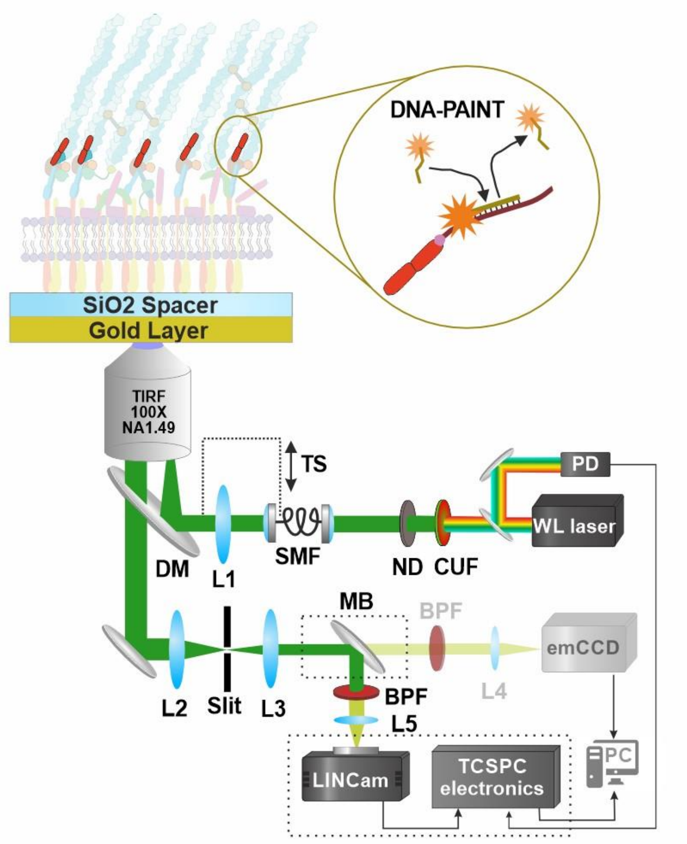


***Figure S1**: Wide-field FLIM optical setup for MIET-PAINT imaging. The setup is equipped with the pulsed super-continuum white light laser, emCCD camera, and with single-molecule sensitive lifetime camera LINCam25 from Photonscore. The setup allows for data acquisition using different illumination schemes: EPI, HILO or TIRF. Sample geometry and DNA-PAINT imaging schematics are shown.*

### Confocal MIET-PAINT

**Optical setup:** Confocal MIET-PAINT measurements were performed using a custom-built confocal microscopy setup described previously [REF to confocal SMLM]; see Figure Sx. Briefly, a white light (WL) laser (Koheras SuperK Power) with a fixed repetition rate of 40 MHz was used. An acousto-optic tunable filter (AOTF) (Koheras SpectraK Dual) attached to the WL laser output provided flexibility in the excitation wavelength. The laser beam was coupled into a single-mode fiber (SMF) (PMC-460Si-3.0-NA012 3APC-150-P, Schäfter+Kirchhoff) using a fiber coupler (60SMS-1-4-RGBV-11-47, Schäfter+Kirchhoff), and was decoupled and collimated using a 10× air objective (UPlanSApo 10×, 0.40 NA, Olympus). A quad-band dichroic mirror (DM) (ZT405/488/561/640rpc, Chroma) was used to reflect the excitation light towards the specimen and also to separate it from the emitted light. The excitation beam passed through a fast laser scanning system (FLIMbee, PicoQuant GmbH). The scanning system deflected the beam to image large regions of interest while maintaining the focus position by directing the laser beam onto the back focal plane of the objective (UApo N 100×, 1.49 NA oil, Olympus). The region of interest and focus plane were controlled by a manual XY stage (Olympus) and a z-piezo stage (Nano-ZL100, MadCityLabs), respectively. The emitted fluorescence light was collected using the same objective and focused onto a pinhole with a 100 µm diameter (PH) (P100S, Thorlabs) using a 180 mm achromatic lens (L1) (AC508-180-AB, Thorlabs). The emission light was then collected and collimated by a 100 mm lens (L2) (AC508-200-A, Thorlabs). Long-pass filters (LP) (488 LP, 647 LP Edge Basic, Semrock) were used to block excitation laser light in the emission path. Band-pass filters (BP) (BrightLine HC 692/40, Semrock) were used to further reject scattered excitation light. Finally, the emission light was focused onto a SPAD detector (SPCM-AQRH, Excelitas) using an achromatic lens (L3) (AC254-030-A-ML, Thorlabs). The output signal from the photon detector was recorded using a TCSPC system (HydraHarp 400, PicoQuant).

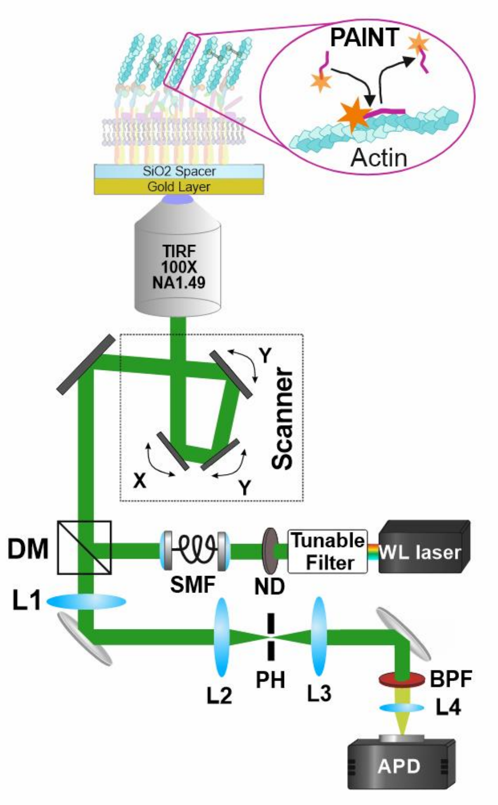


**Figure S2.** Schematic representation of the custom-built confocal SMLM optical setup used for confocal MIET-PAINT measurements. The setup is equipped with white light laser source, fast scanner and TCSPC system as a lifetime modality.

*Experimental conditions and image analysis*

Data was acquired using commercial software from PicoQuant (SymPhoTime 64), which controlled both the TCSPC and scanner systems[2].

For confocal MIET-PAINT measurement, regions of interest measuring 20 × 20 µm² were scanned. A virtual pixel size was set to be 100 nm and a dwell time of 2.5 µs/pixel. A time series of 50.000 frames was recorded. During the analysis, additional time bin of 5 frames was applied.

### Theoretical lifetime-height dependency - MIET curves

The theoretical MIET curves used for lifetime-height conversion for DNA-Atto 550 and Peptide-Cy3B are shown in Figure S3.

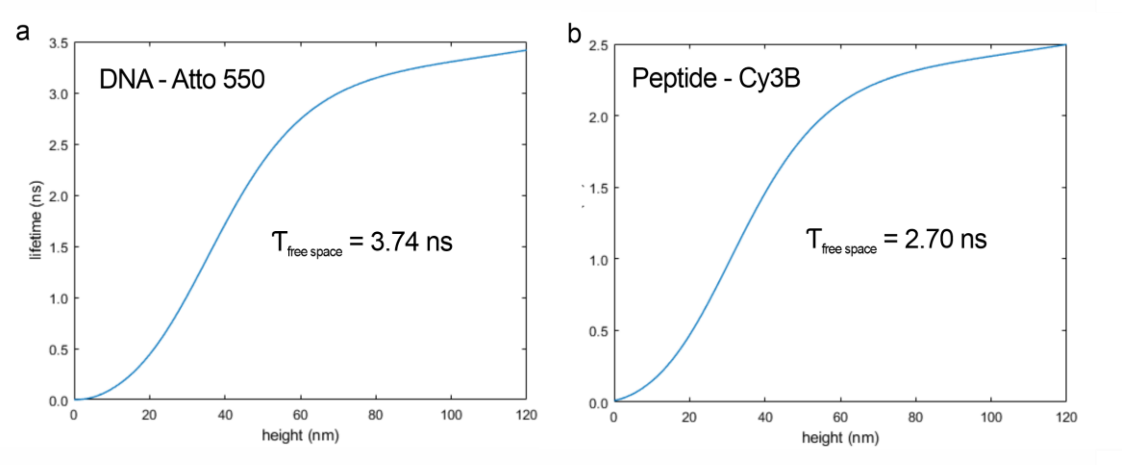


***Figure S3**: Theoretical MIET lifetime-height dependency. MIET curves for DNA-Atto 550 (a) and peptide-Cy3B (b). The values for the free space lifetimes are on top of the plots.*

### Buffer solutions

Following buffer solutions were used: MIET-PAINT validation was performed in imaging buffer containing 10 mM Tris, pH 8.0, 1 mM EDTA, and 500 mM NaCl; MIET PAINT FAC imaging was performed in imaging buffer phosphate-buffered saline (PBS) with 500 mM NaCl.

### Validation of MIET-PAINT

For validation of MIET-PAINT technique and to evaluate the axial localization precision, we performed MIET-PAINT imaging on top of $SiO_2$ spacer with designed thickness. The docking strand was immobilized to $SiO_2$ surface via BSA-biotin neutravidin immobilization strategy. The sample design was as follows: gold coverslip consisted of a thin gold layer of 10 nm and topped with a $SiO_2$ spacer layer of well-defined thickness. We prepared substrates with four different $SiO_2$ layer thicknesses: 20, 30, 40, and 50 nm. The BSA biotin-neutravidin immobilization sandwich added ~12 nm to the total emitter height[3] and was accounted for in the total height estimation. Then, the total heights above the gold layer were 32, 42, 52 and 62 nm. MIET-PAINT imaging was done using the wide-field FLIM optical setup, while the imager-docker binding events were localized and lifetime was extracted for each binding event. Finally, the measured lifetime value was converted into height using MIET theory[4].

***Table S1**: Experimental and calculated lifetime and height values for MIET-PAINT validation experiment.*

| $h_{design}$ (nm) | $\tau_{experiment}$ (ns) | $h_{experiment}$ (nm) | $\Delta h_{CRLB}$ (nm) |
|---|---|---|---|
| 32 | 1.47 ± 0.27 | 36.9 ± 3.9 | 0.6 |
| 42 | 2.07 ± 0.42 | 46.4 ± 6.9 | 1.0 |
| 52 | 2.36 ± 0.14 | 51.5 ± 2.9 | 1.4 |
| 62 | 2.66 ± 0.18 | 58.6 ± 4.1 | 2.02 |
| Glass | 3.62 ± 0.12 | - | - |

### Fluorescence lifetime decay curves

In this section we show the Time Correlated Single Photon Counting (TCSPC) curves for MIET-PAINT data acquired for the different spacers used for validation of MIET-PAINT concept.

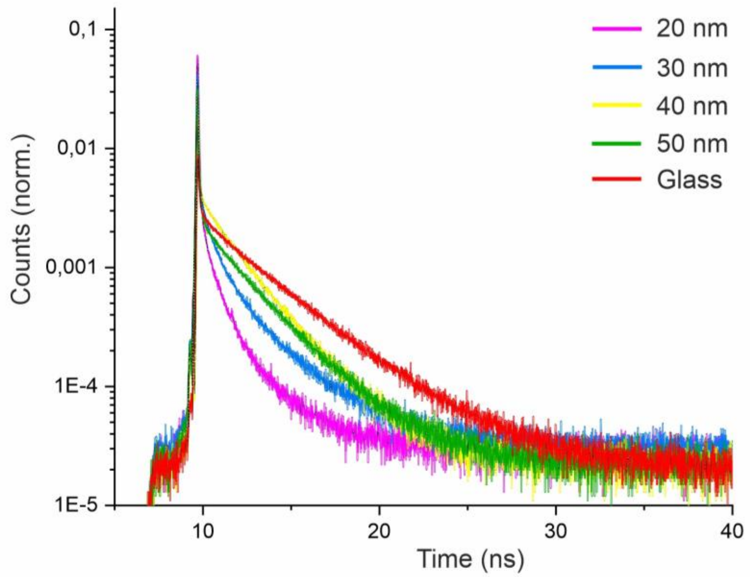


***Figure S4****: TCSPC curves for MIET-PAINT data of different spacers shown in Figure 1 in the main text. All curves were normalized so that area under the curve equals 1.*

### Validation of labeling specificity

In this section we demonstrate the specificity of Zyxin labeling for MIET-PAINT imaging by a comparison between a wide-field GFP image of Zyxin and the super-resolution DNA-PAINT / MIET-PAINT imaging of the same cell[5]. We observe the remarkable correlation between the two images types confirming the specificity of labeling using anti-GFP Nb, as detailed in Methods section.

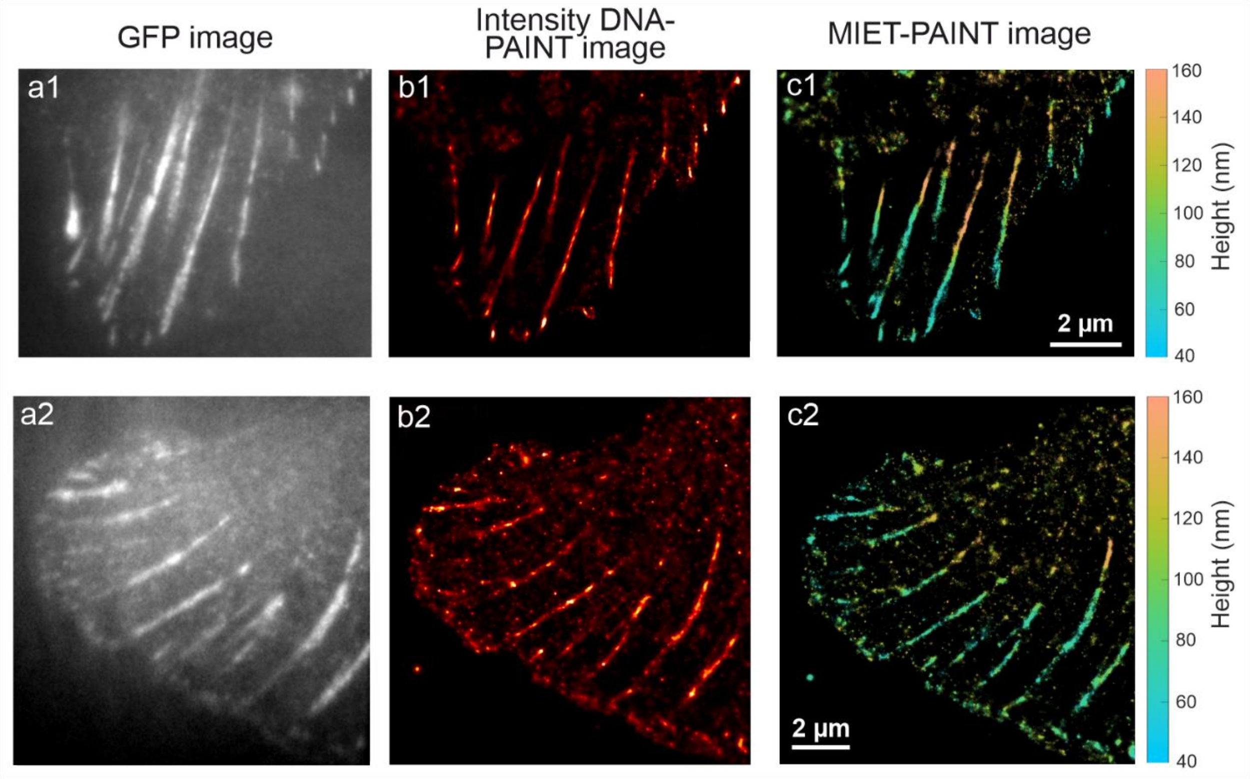


***Figure S5***: *Validation of specificity of Zyxin labeling for cell 1 (main text) and cell 2. (a) Wide-field image of GFP channel, (b) DNA-PAINT and (c) MIET-PAINT images of Zyxin.*

### Additional wide-field MIET-PAINT images

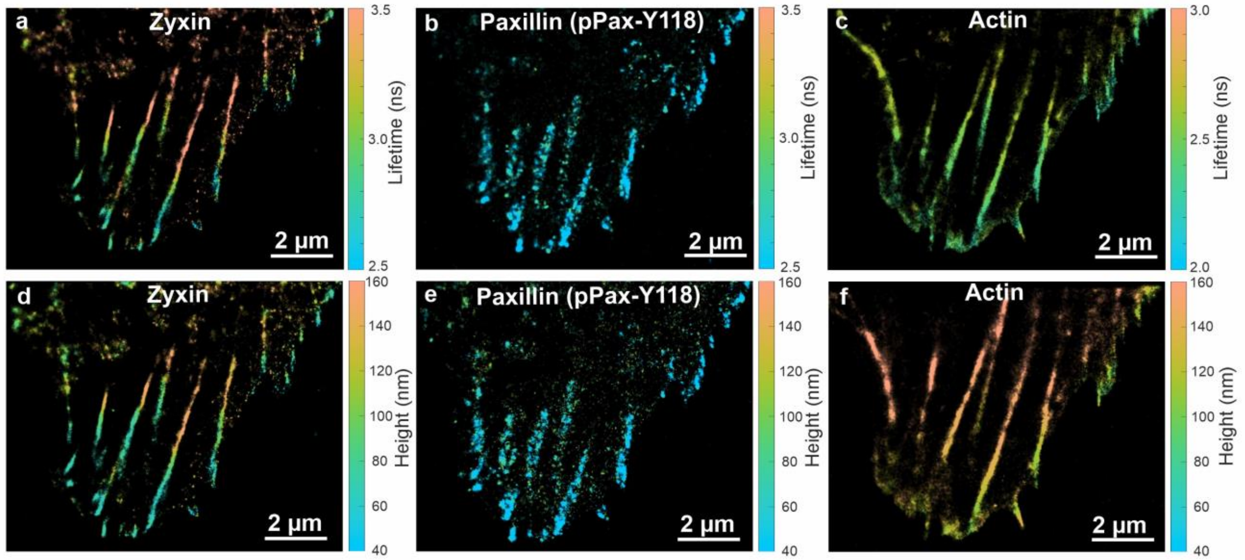


***Figure S6***: *Wide-field MIET-PAINT images of the U2OS cell from Figure 2 (main text). Fluorescence lifetime images of zyxin (a), paxillin (pPax-Y118) (b), and actin (c). Using MIET theory, lifetime images were converted into corresponding height images (d-f). All scale bars are 2 µm. The numbers depict the line height profiles along specific stress fibers*.

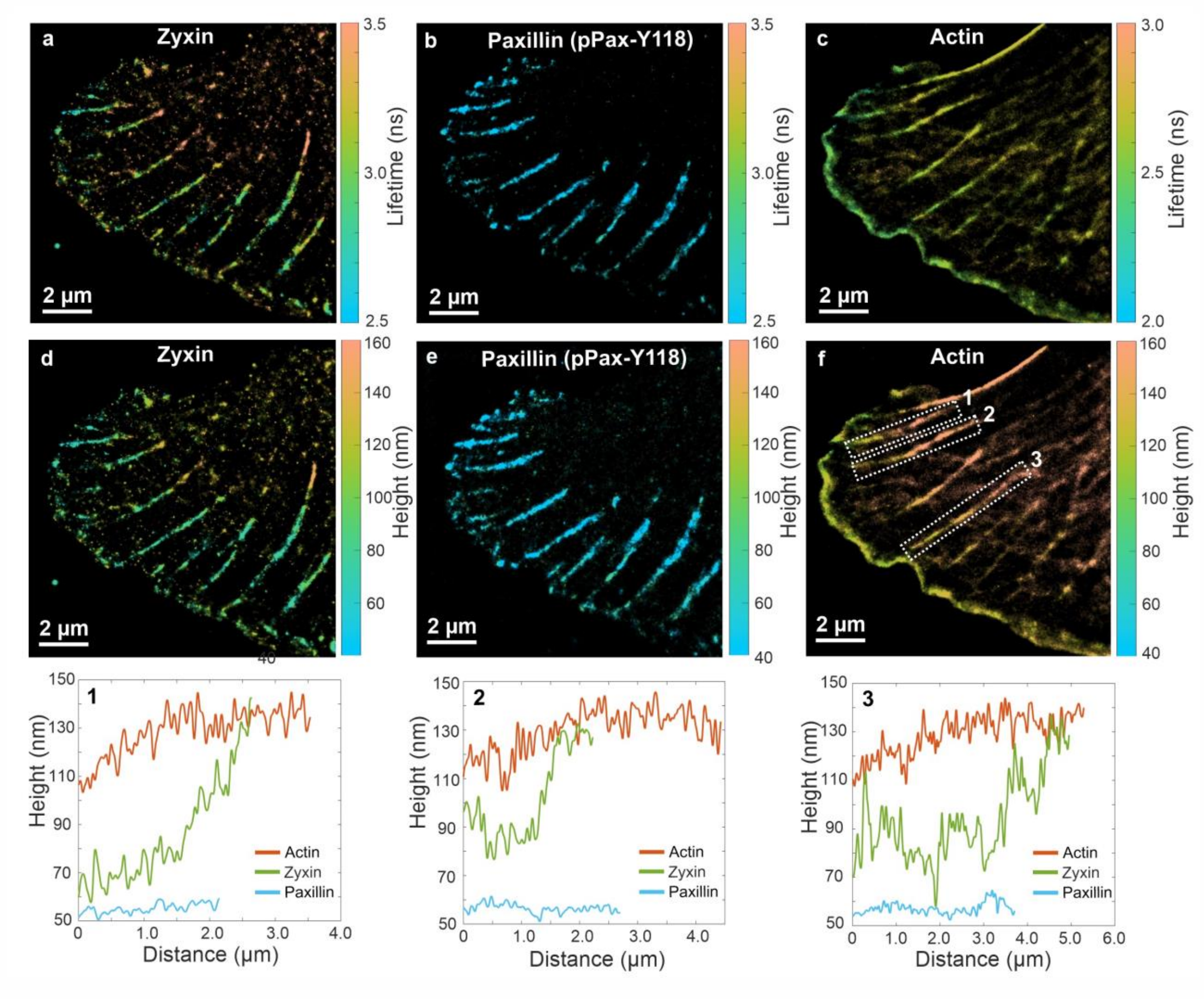


***Figure S7***: *Multiplexed wide-field MIET-PAINT image of U2OS cell #2. Fluorescence lifetime images of zyxin (a), paxillin (pPax-Y118) (b), and actin (c) of the same cell were acquired sequentially. Using MIET theory, lifetime images were converted into corresponding height images (d-f). All scale bars are 2 µm. (g) Regions that were selected to show height profiles of zyxin, paxillin, and actin along the stress fibers as marked in dashed white rectangle in (f). The numbers depict the line height profiles along specific stress fibers.*

### 3D representation of MIET-PAINT images

Here, we rendered a 3D representation of localizations for wide-field MIET-PAINT data from Figure 2 in the main text:

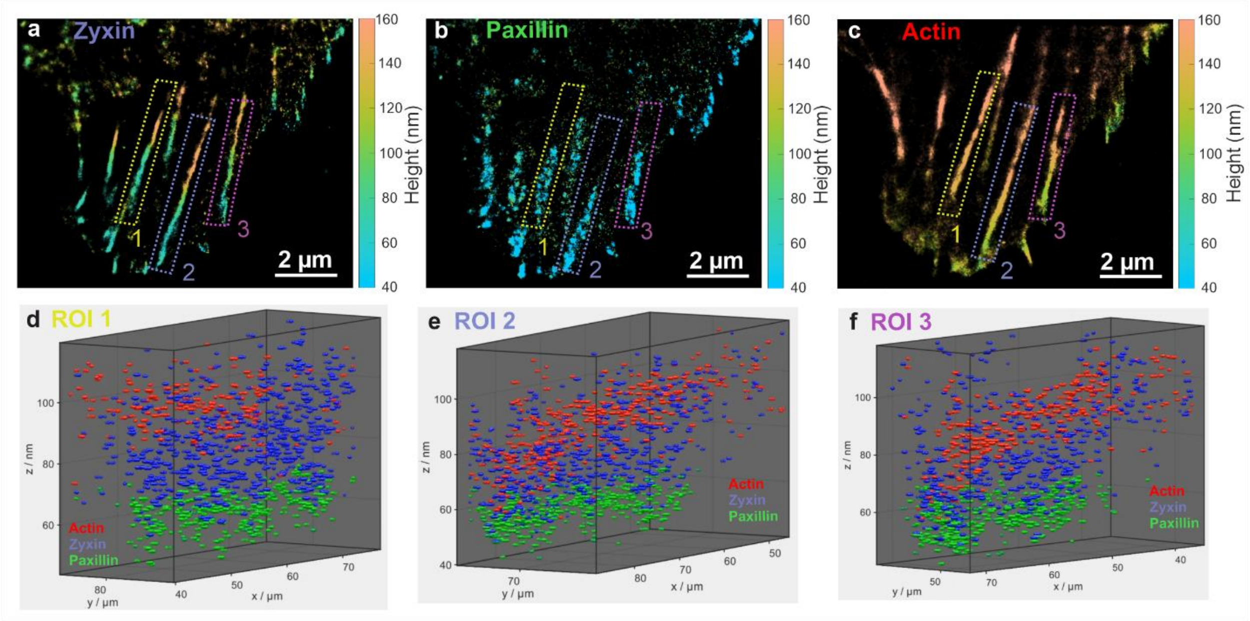


***Figure S8***: *3D Representation of MIET-PAINT data for FAC proteins. The 3D arrangements for Zyxin (a) Paxillin (b) and Actin (c) from Figure 2 in the main text for ROI 1-3 (d-f).*

### Additional confocal MIET-PAINT images

Additional lifetime confocal MIET-PAINT images for the cell shown in Figure 3 in the main text.

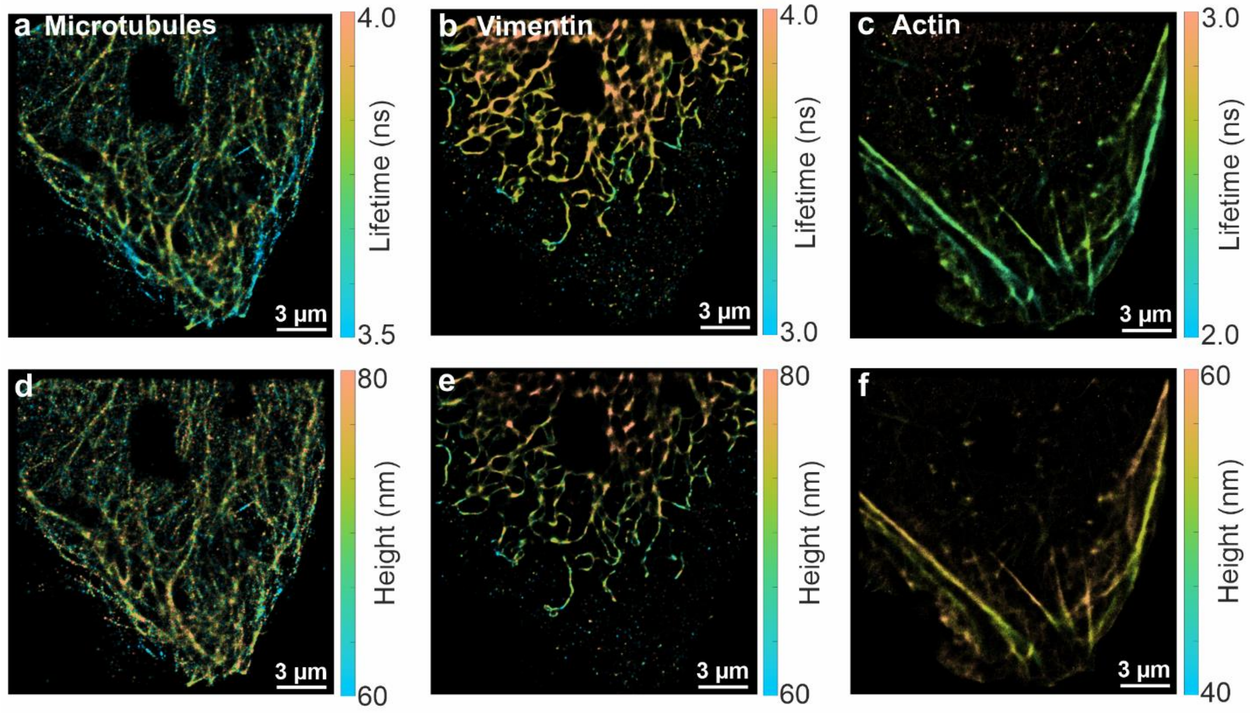


***Figure S9***: *Confocal MIET-PAINT images of cytoskeleton in U2OS cell. Fluorescence lifetime images of microtubules (a), vimentin (b), and actin (c) of the same cell. MIET curve was used to convert lifetime value for each localization into height, resulting in a correspondent height MIET-PAINT images (d-f). All scale bars are 3 µm.*

Lifetime and height confocal MIET-PAINT images of the cell #2.

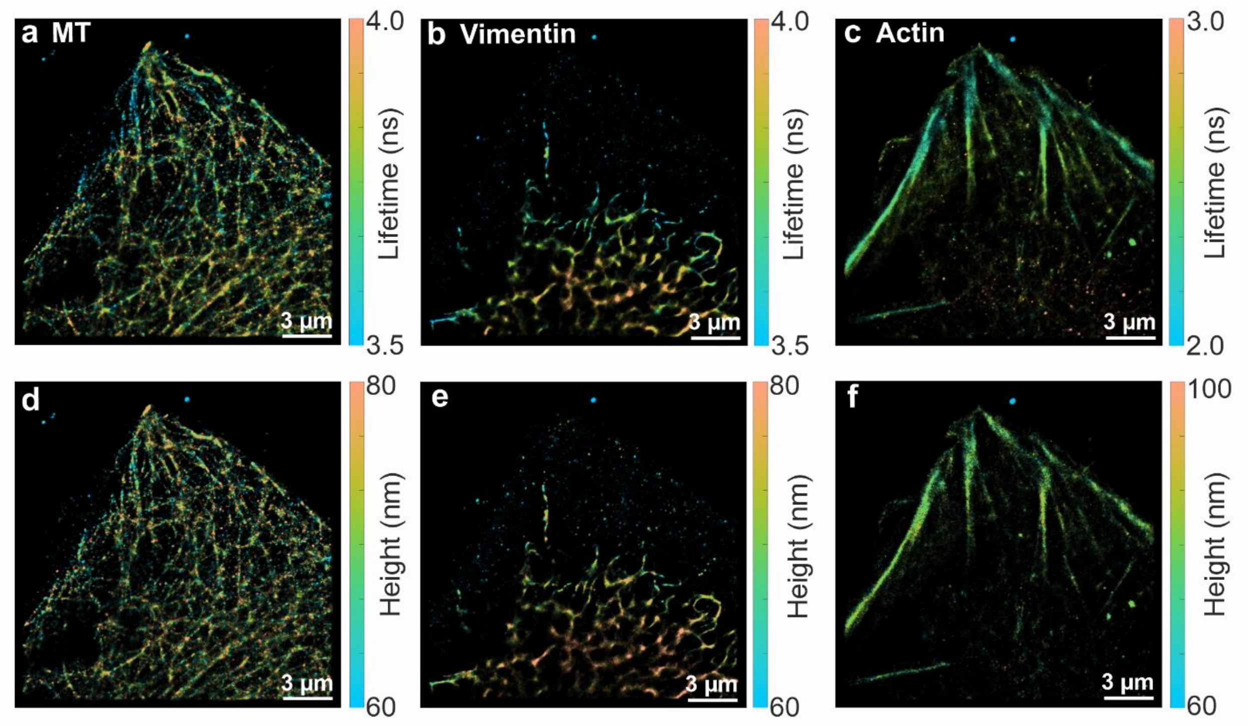


***Figure S10**: Confocal MIET-PAINT images of cytoskeleton in U2OS cell. Fluorescence lifetime images of microtubules (a), vimentin (b), and actin (c) of the cell #2. MIET curve was used to convert lifetime value for each localization into height, resulting in a correspondent height MIET-PAINT images (d-f). All scale bars are 3 µm.*

Lifetime and height confocal MIET-PAINT images of the cell #3.

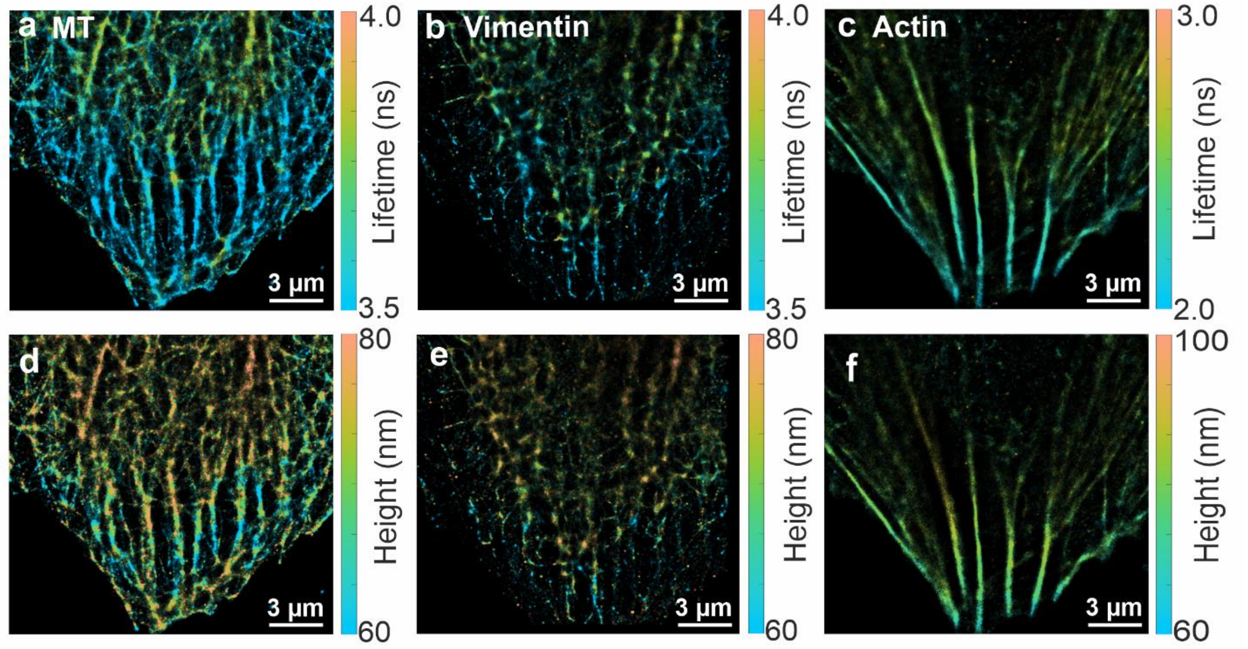


***Figure S11***: *Confocal MIET-PAINT images of cytoskeleton in U2OS cell. Fluorescence lifetime images of microtubules (a), vimentin (b), and actin (c) of the cell #3. MIET curve was used to convert lifetime value for each localization into height, resulting in a correspondent height MIET-PAINT images (d-f). All scale bars are 3 µm.*

### Quantitative description of MIET-PAINT images

In this section, we provide a quantitative assessment of wide-field and confocal MIET-PAINT datasets corresponding to the cell images shown in Figures 2 and 3 of the main text, as well as SI Figures S7, S9–S11. Specifically, we report the average lateral and axial localization precisions obtained for all imaged targets. Lateral localization precision was estimated using a modified Mortensen's equation[6,7]. Axial localization precision was directly derived from the MIET-PAINT data by calculating the Cramér–Rao lower bound (CRLB) of the fluorescence lifetime for each localization, based on the detected photon counts[8]. The resulting lifetime uncertainties were then converted into axial uncertainties via the MIET calibration curve, and the mean precision values were computed for each dataset.

***Table S2***: *Lateral and axial localization precision of wide-field MIET-PAINT images.*

| Figure / target | Average lateral loc. precision (nm) | Average axial loc. precision CRLB (nm) |
|---|---|---|
| Fig. 2 / Zyxin | 14.6 | 3,9 |
| Fig. 2 / Paxillin | 21.6 | 3,9 |
| Fig. 2 / Actin | 25.8 | 8,4 |
| Fig. S7 / Zyxin | 21.4 | 4,7 |
| Fig. S7 / Paxillin | 25.9 | 3,2 |
| Fig. S7 / Actin | 23.6 | 7,5 |

***Table S3**: Lateral and axial localization precision of confocal MIET-PAINT images.*

| Figure / target | Average lateral loc. precision (nm) | Average axial loc. precision CRLB (nm) |
|---|---|---|
| Fig. 3/S9 / Microtubules | 10.8 | 3,6 |
| Fig. 3/S9 / Vimentin | 13.7 | 2,8 |
| Fig. 3/S9 / Actin | 14.3 | 4,9 |
| Fig. S10 / Microtubules | 9.5 | 2,7 |
| Fig. S10 / Vimentin | 15.2 | 2,5 |
| Fig. S10 / Actin | 13.0 | 4,7 |
| Fig. S11 / Microtubules | 10.5 | 2,7 |
| Fig. S11 / Vimentin | 11.8 | 2,7 |
| Fig. S11 / Actin | 14.9 | 4,7 |

### DNA sequences of imager and docker strands

***Table S4**: Detailed DNA sequences and modification of imagers and dockers strands.*

| name | sequence 5′ → 3′ | 5′-modification | 3′-modification |
|---|---|---|---|
| imager R1 | AGGAGGATT | - | Atto 550 |
| imager R4 | TGTGTGTTT | - | Atto 550 |
| imager R3 | GAGAGAGAAA | - | Atto 550 |
| imager R6 | TTGTTGTTT | - | Atto 550 |
| docker R1* | TCCTCCTCCTCCTCCTCCT | Azide | - |
| docker R4* | ACACACACACACACACACA | Azide | - |
| docker R3* | CTCTCTCTCTCTCTCTCTC | Azide | - |
| docker R6* | AACAACAACAACAACAACAA | Azide | - |

### Targets correlation analysis – Pearson coefficient

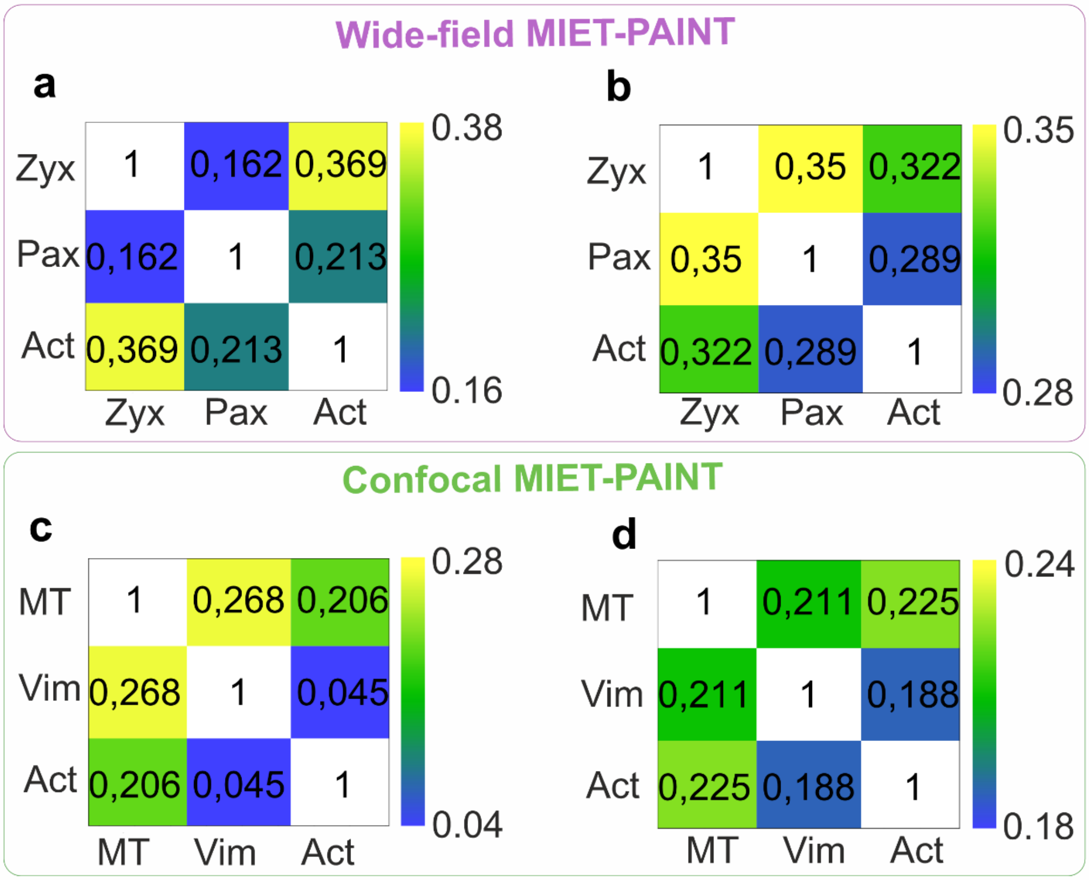


***Figure S12***: *Heatmaps with Pearson coefficient estimate for targets cross-correlation analysis for wide-field MIET-PAINT cell 1 (a) from the main text and cell 2 (b) from the SI; and confocal MIET-PAINT cell 2 (c) and cell 3 (d) from the SI.*
